# Deep learning for rapid and reproducible histology scoring of lung injury in a porcine model

**DOI:** 10.1101/2023.05.12.540340

**Authors:** Iran A. N. Silva, Salma Kazemi Rashed, Ludwig Hedlund, August Lidfeldt, Nika Gvazava, John Stegmayr, Valeriia Skoryk, Sonja Aits, Darcy E Wagner

**Affiliations:** Lung Bioengineering and Regeneration, Department of Experimental Medical Sciences, Faculty of Medicine, Lund University, Lund, Sweden; Wallenberg Center for Molecular Medicine, Faculty of Medicine, Lund University, Lund, Sweden; Stem Cell Center, Faculty of Medicine, Lund University, Lund, Sweden; NanoLund, Lund University, Lund, Sweden; Cell Death, Lysosomes and Artificial Intelligence Group, Department of Experimental Medical Science, Faculty of Medicine, Lund University, Lund, Sweden; Medical Microspectroscopy, Department of Experimental Medical Science, Faculty of Medicine, Lund University, Lund, Sweden; Lund University Profile Area Artificial and Natural Cognition, Lund, Sweden; Lund University Profile Area Nature-Based Future Solutions, Lund, Sweden; Lund University Cancer Centre, Lund, Sweden

**Keywords:** Score System, Deep Learning, Computer vision, Convolutional neural network, Acute Lung Injury, Acute respiratory distress syndrome, Histology

## Abstract

Acute respiratory distress syndrome (ARDS) is a life-threatening condition with mortality rates between 30-50%. Although *in vitro* models replicate some aspects of ARDS, small and large animal models remain the primary research tools due to the multifactorial nature of the disease. When using these animal models, histology serves as the gold standard method to confirm lung injury and exclude other diagnoses as high-resolution chest images are often not feasible. Semi-quantitative scoring performed by independent observers is the most common form of histologic analysis in pre-clinical animal models of ARDS. Despite progress in standardizing analysis procedures, objectively comparing histological injuries remains challenging, even for highly-trained pathologists. Standardized scoring simplifies the task and allows better comparisons between research groups and across different injury models, but it is time-consuming, and interobserver variability remains a significant concern. Convolutional neural networks (CNNs), which have emerged as a key tool in image analysis, could automate this process, potentially enabling faster and more reproducible analysis. Here we explored the reproducibility of human standardized scoring for an animal model of ARDS and its suitability for training CNNs for automated scoring at the whole slide level. We found large variations between human scorers, even for pre-clinical experts and board-certified pathologies in evaluating ARDS animal models. We demonstrate that CNNs (VGG16, EfficientNetB4) are suitable for automated scoring and achieve up to 83% F1-score and 78% accuracy. Thus, CNNs for histopathological classification of acute lung injury could help reduce human variability and eliminate a time-consuming manual research task with acceptable performance.

## 1. Introduction

Acute respiratory distress syndrome (ARDS) is a highly heterogenous and life-threatening disease with mortality rates of around 30-50% worldwide (1–3). Depending on the source of the injury (*e.g.* specific pathogen, trauma, *etc.*), patients present with different local or systemic pathophysiological features (4). While *in vitro*, for example cell culture-based, models can recapitulate some features of ARDS, the disease is multifactorial and therefore small and large animal models remain the main preclinical research tools. Both rodent and pig models are commonly used, but the choice of small or large animal model depends on the disease features to be modelled or the type of therapy to be evaluated (5, 6).

The clinical diagnosis of ARDS in patients is based on the Berlin criteria which encompasses 4 main characteristics (7, 8): known timing of injury within one week, bilateral opacities observed via chest imaging (X-ray or computed tomography (CT) scan) which exclude atelectasis or lung nodules, exclusion of edema from cardiac failure, and compromised oxygenation (divided into three levels: mild, moderate or severe). Although animal models can also be assessed using the Berlin criteria, chest imaging with CT can be difficult in practice due to the short experimental duration in injury models for ARDS, scarcity of equipment, or difficulties with the logistics of transporting animals (9, 10). In the absence of high-resolution chest images, histological assessment instead serves as a gold standard method to exclude other differential diagnoses and is similarly used to determine the level of injury in animal models (6, 11, 12). Semi-quantitative scoring of damage features by human observers is the most common analysis form used to assess histology samples from pre-clinical animal models of ARDS (13, 14). The most widely used score system was originally developed to score acute lung injury in mice, and has been applied to large animal models in a few studies (9). However, interspecies variation in morphological and physiological features, for example measurements of arterial blood gases, are feasible in large animals, but are challenging to implement in small animal models, such as mice, and these challenges can lead to misinterpretation of findings and affect the validity of the scoring system (9, 10). Another limitation of human based evaluation of histological slides is the fact that visual assessment, even with standardized and improved scoring systems, carries a certain degree of subjectivity (15). Intra- and inter-observer variability therefore limits reproducibility, even among highly trained board-certified pathologists (16, 17). This ultimately impacts the sensitivity with which semi-quantitative scoring methods can detect meaningful biological differences in histological samples. Technologies which accurately automate this task at the level of expert human observers could thus not only decrease the analysis time but could additionally reduce differences which arise due to variability between human observers (especially those with different levels of experience) and improve the sensitivity at which histological samples can be assessed (10, 18–23).

Artificial intelligence tools, convolutional neural networks (CNNs) in particular, have gained attention for their use in a wide range of image analysis applications, including medical research and clinical diagnostics. In relation to lung disease, CNNs have for example been used for classification of lung tumor types, including both histological and genetic phenotypes (24–27), murine lung fibrosis (28) and for segmentation of features of interest for diagnosis of pulmonary disease, treatment or follow-up (29–31). The use of CNNs not only greatly reduces analysis time and cost, but also the bias and reproducibility issues displayed by human observers. In many cases, CNNs also surpass human observers in image analysis accuracy because the models more readily detect subtle changes and variations that humans find difficult to see (32). Existing CNN-based histological image analysis has focused on characterizing clinical samples (24, 27, 30) or histological remodeling in murine models of chronic lung disease (28) and thus far there have been no attempts to characterize animal models of ARDS. A CNN-based imaging tool would be a major advancement in particular for large animal models, as multiple samples are required to reduce sampling bias, making the analysis especially time-consuming and expensive.

Here we assessed whether our recently developed system for semi-quantitative scoring of histological lung injury in a porcine model of ARDS could be suitable for knowledge transfer to CNNs (14). As a proof-of-concept, we then trained CNNs to classify tissue damage as mild, medium, or high, finding them able to perform similarly or better than typical human scorers.

## 2. Methods

### 2.1 Image data for manual, human-based scoring

We used images of digitally scanned hematoxylin and eosin (H&E) stained histological slides which had been generated from a porcine model of lipopolysaccharide (LPS) acute lung injury and previously deposited in the Biostudies archive of the European Bioinformatics Institute (EMBL-EBI) (https://www.ebi.ac.uk/biostudies/studies/S-BIAD419). Experimental groups and the accompanying total injury scores, which were described in the original publications (13, 14) and annotated in the metadata of S-BIAD419, included control (baseline samples), lipopolysaccharide (LPS) administration, mechanical ventilation (MV), extracorporeal membrane oxygenation (ECMO) or combinations of these (**Supplemental Figure 1 and 2a)**. As detailed in the original publications describing the animal model, all animals who received LPS administration fulfilled the Berlin criteria of ARDS (33) by two serial blood gas measurements prior to being stratified into MV only or MV and ECMO experimental groups (13, 14). Experimental procedures and data collection for histological image collection were described in the previous publications and S-BIAD419 (5, 13, 14). In short, bright field images of H&E-stained histological slides were obtained using a VS120 virtual microscopy slide scanning system (Olympus, Tokyo, Japan). Acquisition settings were the same for all slides. Each virtual slide contained several sections from a single animal and experimental time point. Representative images for each slide were then cut out computationally from different locations on the slide, starting from the center of the biopsy and avoiding airways, at three different digital magnifications (4x, 10x and 20x) in the OlyVIA 3.8 Olympus Software viewer. Settings during image extraction were the same for all virtual slides: Brightness: 50%, Contrast: 50% and Gamma: 1. Extracted images were assembled in a Microsoft PowerPoint file for scoring (images from one slide per page and containing the 3 different magnifications). Observers were given 49 virtual histological slides to score. This included 45 unique slides as well as 4 duplicate slides which were randomly inserted without informing observers. These repeat slides were utilized in S-BIAD419 as an internal quality control of self-agreement for each observer and to prevent observers who knew the study design from allocating pigs to known experimental groups. Each observer assigned one score per slide for each feature and a total score per slide was assigned by summing up the total of the seven individual feature scores.

### 2.2 Independent validation of the previously described scoring system

To further explore the inter-observer reproducibility of our previously described scoring system, we asked 5 new observers (validation cohort) to repeat the scoring as previously described (13, 14) in the current work. Additionally, a subset of these observers was asked to repeat the same scoring after at least 1 month to test intra-observer reproducibility (**Figure 1a and Supplemental Figure 1a).** Briefly, the score system consists of 7 features that reflect damage features we and others have previously observed in the LPS porcine model of ARDS: inflammatory cells, hyaline membranes, proteinaceous debris, thickening of alveolar walls, hemorrhage, atelectasis and a general assessment of enhanced injury. Observers were instructed to provide a score in the range between 0 (no damage) and 8 (extensive damage), resulting in a total score in the range of 0 to 56. The initial scoring results deposited with the imaging metadata from our previous publications (S-BIAD419; referred to as cohort “A”) was performed by six reviewers with different levels of expertise: 2 novices (N), 2 moderates (M), and 2 pre-clinical experts (pE). These reviewers were named N1, N2, M1, M2, pE1 and pE2 (**Figure 1a, Supplemental Figure 1**). The validation cohort (“B”) contained 5 reviewers (M3, M4, pE3 and pE4, **Figure 1a, Supplemental Figure 1**). To assess the self-agreement of the reviewers, three reviewers scored images again after at least 1 month (M3, pE3 and pE4) as a repetition cohort. In addition, two board-certified clinical pathologists (bcP1 and bcP2) scored the images. For comparison with other score systems, scores ranging from 0-56 were then scaled to be on a range of 0-100: Normalized total score = (total score/56) x 100.

**Figure 1.**
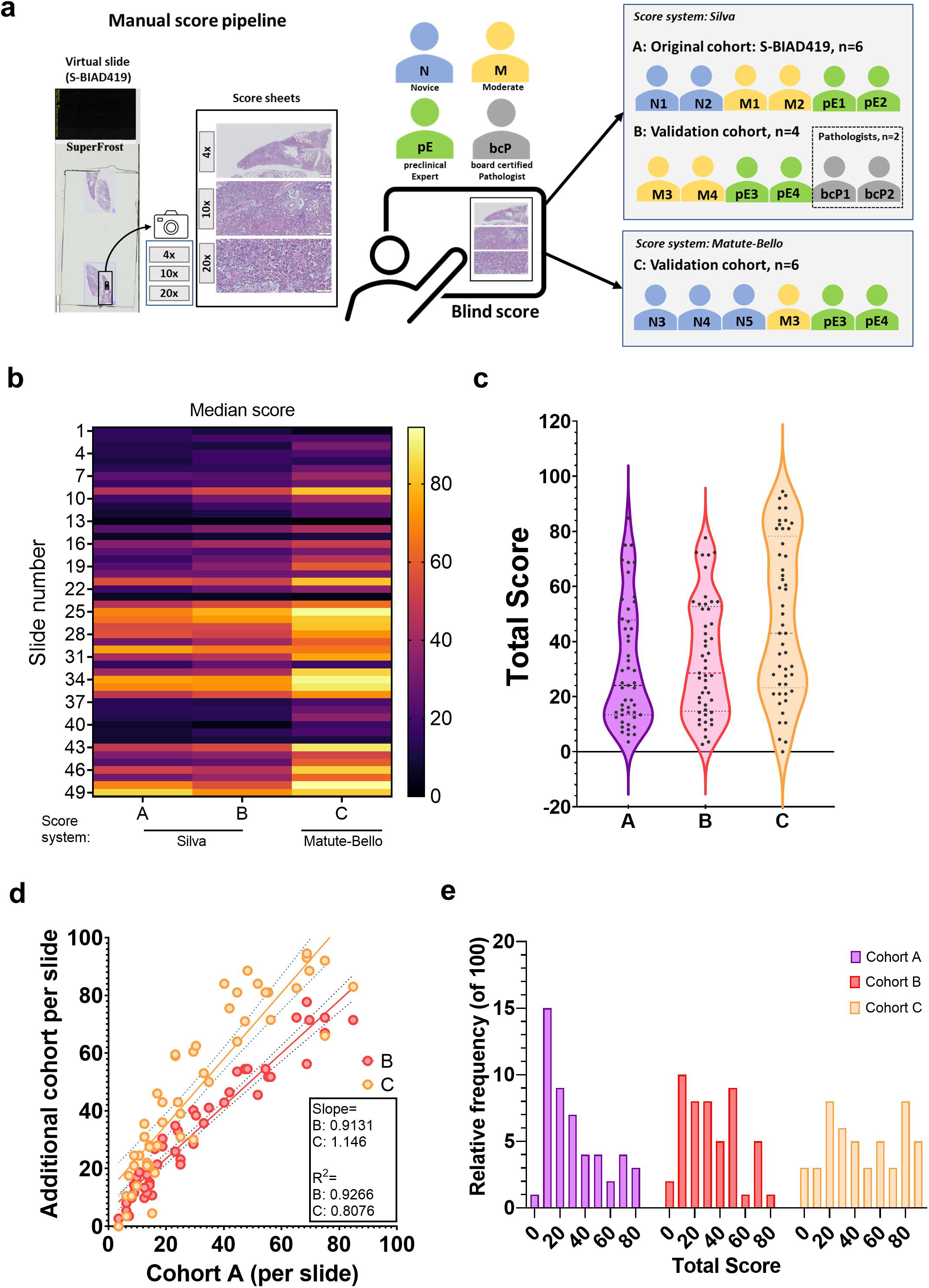
A recently developed semi-quantitative score system for acute lung injury is suited to generate ground truth labels for training deep neural networks. **(A)** Schematics of the manual scoring pipeline. The lettering signifying the original (A), validation (B) and Matute-Bello (C) cohorts is used throughout the other figure panels **(B)** Heatmap of median total score per slide for the selected cohorts ordered by experimental groups: control, ECMO, ECMO+LPS, MV and MV+LPS groups. Scores were normalized to be within a range of 0-100 for this figure and the same slide order was used in all subsequent figure panels. **(C)** Violin plot comparing the distributions of the normalized median total scores from the three cohorts. **(D)** Scatter plot comparing the normalized median total scores of the two new cohorts with the original *Silva et al.* cohort. N refers to the number of scorers. Slope refers to the slope of the regression line. **(E)** Histogram demonstrating the distribution of the normalized median total scores of the selected cohorts.

To compare our scoring system to the current widely-used score system for small animal models, which was originally described by Matute-Bello *et al.* (9) and reaffirmed in a recent workshop report by a group of experts in pre-clinical models of ARDS (10), we had a subset of observers perform additional scoring using a version of the Matute-Bello score system which we modified for use with virtual histological slides. The score system described by Matute-Bello slightly differs from the system that we developed for our large animal model and includes scoring of 5 features: A: Neutrophils in the alveolar space, B: Neutrophils in the interstitial space, C: Hyaline membranes, D: Proteinaceous debris filling the airspace and E: Alveolar septal thickening. Because it would be extremely time-consuming to perform scoring of so many slides and with multiple observers using high magnification regions (i.e. 400x total magnification), we asked observers to score at the slide-level similar to our previously described scoring system. To avoid the potential of introducing bias by using other areas of the slides, we used the same PowerPoint sheets as previously generated for all observers and score systems. For the modified Matute-Bello system, we asked observers to assign a score of 0, 1, or 2 for each feature (labeled A-E in the following formula) which represented the most common extent of injury they observed in the images. The total score was then calculated using the recommended weighted formula (9): Score = [(20 × A) + (14 × B) + (7 × C) + (7 × D) + (2 × E)]/ (number of fields × 100). This yields a value between 0-1. Therefore, for comparison, scores were normalized to be in a range of 0-100 by multiplying the total score with 100. This scoring was performed by six observers (N3, N4, N5, M3, pE3 and pE4) and the data referred to as the “C” cohort.

### 2.3 Statistical analysis and data visualization

Aggregated scoring results are shown as median with standard deviation as error bars in plots generated in GraphPad Prism 9 (GraphPad Software Inc, La Jolla, CA, USA). Median total scores for each slide were compared between cohorts using simple linear regression. To assess the reliability or internal consistency of the score system, the Spearman rank correlation was calculated between all reviewers and across all cohorts. The range of scores per slide were calculated for each expertise level and for each cohort. Linear regression analysis was performed to assess the intra-observer variation. P-values <0.05 indicate a significant difference. Plots related to the deep learning pipeline were generated with Matplotlib (version 3.5.9) and seaborn (version 0.11.2) python packages.

### 2.4 Dataset pre-processing and augmentation for training of deep neural networks

For the supervised training of CNNs, up to 10 non-overlapping tissue areas were collected per slide and corresponding labels from 5 observers (N1, M1, M2, pE1 and pE2) were used (13) (**Supplemental Figure 1**). Images were extracted from the digital whole-slide images by manual selection from the OlyVIA 3.8 Olympus Software viewer in order to mimic how human observers experienced the images digitally (*i.e.* stitched montages of individually acquired images). Extracted images were then saved as 8-bit RGB Tif images with pixel dimensions of 890 x 1674 or 586 x 1335 or 817 x 1635 as automatically set by the software. For training of the deep neural networks, three of the duplicate slides were excluded and one was excluded due to technical problems with the viewer software. As described above, observers had quantified the seven damage features using a score range of 0 to 8, resulting in a total range of 0 (no damage) to 56 (most severe damage in all features) for the total score of each slide. The median of the total score from the five observers were converted to class labels, which reflected different levels of damage and served as ground truth labels for the CNNs. Ground truth labels were assigned based on a trimodal distribution, representing low, medium and high histological damage. A trimodal distribution was selected based on the anticipated and potential heterogeneity of histological injury existing within a slide and was selected after evaluating the distribution of scores assigned to the tiles in our dataset. Ranges were constructed so that there was at least one major peak in each class and that each ground truth label had approximately similar number of tiles.

To match the expected input size of the CNNs and increase the number of training examples fed to the networks, each image was cut into adjacent non-overlapping tiles with a size of 224 x 224 pixels, starting from the top left corner. The number of tiles cropped from each image was dependent on the original image size (**Supplemental Figure 2a-b**). Only tiles that had image segments covering the entire tile were retained and smaller, marginal tiles, which also included the areas containing text and size bars, were discarded (**Supplemental Figure 2b**).

As the dataset was deemed too small to set aside a separate test set for CNN evaluation, the dataset was instead prepared for 3-fold cross-validation (34). For this, the data was divided into 3 parts with all the sample groups included in each part (**Supplemental Figure 2c**). All tiles from the same slide were always kept together either in the training or validation set to avoid data leakage. For each fold, two of the parts were then combined for training and the third used as test set to evaluate the trained neural network.

To increase the size of the training data and the robustness of the CNNs towards variation in brightness and blurriness that can be caused by the slide scanning microscope, the training data was augmented for some models in one of three ways (**Supplemental Figure 3a, Supplemental Table 1**): 1. For models indicated with the suffix “_b”, brightness was altered with 50% probability by multiplying pixel intensity values with a randomly chosen value between 0.8 and 1.2 and then clipping the pixel values to the normal 8-bit greyscale range [0,255], and blurring was applied afterwards with 50% probability with kernel sizes of 5 x 5 and horizontal flipping with 50% probability. For this, a custom algorithm was written. 2. For models indicated with the suffix “_r_a”, the Keras (version 2.8.0) (35) “ImageDataGenerator” function was applied with rotation in the range of –20 to +20 degrees to mimic slight rotations as well as width and height shifting, flipping and change of brightness in the range of (0.8 to 1.3). For shifting and rotation the filling mode of empty border pixels formed by the shift or rotation, was set to ‘nearest’ pixels. 3. For models indicated with the suffix “_m_a”, the “ImageDataGenerator” function was applied as for the “_r_a” models for flipping and brightness change, but without shifting and with rotation of 90, 180 and 270 degrees (with a custom written rotation function added to the ImageDataGenerator). Augmentation scripts are deposited at https://github.com/Aitslab/lunghisto.

Images were stored and pre-processed on the LUNARC, Berzelius, Kebnekaise and Alvis high performance computing clusters, which are part of the Swedish National Infrastructure for Computing (SNIC).

### 2.5 Training of deep neural networks

Multiple CNN models were trained to classify the amount of damage as low, medium, or high using the TensorFlow (version 2.8.0) python package with or without the augmentation described above (36). For each training round with a specific model type and input, 3-fold cross validation was performed, meaning that two parts of the data were combined into a set for training and the third used as test set. This was repeated until all three combination possibilities had been used. CNNs were trained from scratch or with transfer learning. Two CNN architectures were compared (**Supplemental Figure 3b-c**): 1. VGG16 backbone (37) + 2 fully connected layers with 128 nodes + Dropout layer with rate of 0.5 + SoftMax layer with 3 class (3,228,291 trainable parameters), and EfficientNetB4 (38) backbone + 2 fully connected layers with 128 nodes + Dropout layer with rate of 0.5 + SoftMax layer with 3 class (11,256,451 trainable parameters). For transfer learning CNNs were initialized with weights pre-trained on ImageNET (39), which are included in the TensorFlow package, and then only the last two fully connected layers and the SoftMax layer were trained whereas the other layers were kept frozen. Training was performed over 25 epochs using the Adam optimizer (learning_rate=0.001). The Keras (35) callbacks function, included in TensorFlow, was used to save the models from those 25 epochs and the best model was chosen based on the tile level accuracy in the respective test set (the part of the data not used for training in the respective fold). The best model was then evaluated fully (see below). CNNs were trained on NVIDIA V100 and A100-SXM4-40GB GPU nodes of the Alvis and Berzelius high-performance clusters of the Swedish National Infrastructure for Computing (SNIC).

### 2.6 Evaluation of trained deep neural networks

Each trained CNN model was evaluated by making predictions on all tiles from the respective test set (the part of the data not used for training in the respective fold) and comparing the prediction to the ground truth labels. To obtain slide label predictions, the predictions for all tiles from the respective slide were aggregated by majority vote. We generated 3 x 3 confusion matrices, which display the correct vs the predicted tile labels for all three damage classes and calculated a range of informative metrics on the slide level with the Keras (version 2.8.0) and scikit-learn (v. 1.0.1) python libraries. The following metrics were used for evaluating the individual label classes: 1. Precision: number of tiles correctly predicted to be in a class (true positives) divided by the number of all tiles predicted to be in this class; 2. Recall: number of tiles correctly predicted to be in a class (true positives) divided by total number of tiles in this class; 3. F1-score: harmonic average of recall and precision = 2 x precision x recall/(precision + recall). The following metrics were used for evaluation across classes to account for the imbalance between classes: 1) Micro-average of precision, recall, and F1-score, which is equivalent to overall accuracy. 2) Weighted average precision and recall: the sum of the corresponding class metrics, each multiplied with the fraction of tiles that belong to the respective class. 3) weighted average F1-score: harmonic mean of the weighted average precision and weighted average recall. For all metrics, the average across the three folds was then calculated. To obtain confusion matrices that represent all three folds, the number of tiles from the individual matrices from each of the three folds were added.

### 2.7 Ensemble models

For ensemble prediction, the predictions of the three best CNN models were aggregated by the ‘majority vote’ rule on tile and slide level. In the tile level aggregation, all tile labels predicted for a slide by the three best CNN models were pooled together and the decision for the final slide label based on a “majority vote”. In the case of equal number of votes for more than one class, the best CNN model provides the tiebreaker vote (E_m_a). In slide level aggregation, the class given by each CNN model was calculated based on the major class of tiles predicted by that CNN model. These individual predictions were then aggregated into the final slide prediction.

## 3. Results

### 3.1 A recently developed semi-quantitative score system for acute lung injury is suited to generate ground truth for training deep neural networks

To train deep learning models such as CNNs to detect the extent of histological lung injury in a supervised manner, high quality ground-truth labels are needed. For this, images must be annotated using categorical or quantitative values. As semi-quantitative systems are typically used for assessing the extent of histological damage in acute lung injury, we explored whether such systems are suitable for generating ground-truth labels. We examined the robustness and potential suitability of two different scoring systems on the same set of histological slides using multiple, independent observers representing the range of expertise typically available in a pre-clinical lab (**Supplemental Figure 1**). The system we evaluated was the widely used scoring system originally published by Matute-Bello *et al.* as part of an American Thoracic Society Workshop Report of experts in pre-clinical models of ARDS in 2011 (9), which has recently been reinforced as state-of-the-art in the pre-clinical field by a group of pre-clinical experts in acute lung injury animal models (10). It uses a range of 0-2 across 5 damage features and combines them into a total score using a weighted formula. The second scoring system (Silva *et al.* system) was recently developed by our group and uses a range of 0-8 across 7 features and then sums the feature scores to obtain the total score (5, 10, 13, 14).

We used images from 45 slides of H&E-stained lung histology sections which we had digitally scanned and previously deposited in a public repository S-BIAD419 together with Silva *et al.* system scores from 6 independent scorers (termed “A” cohort). These tissue sections were from a porcine model of ARDS with five experimental groups which exhibited different degrees of lung damage: control (baseline), mechanical ventilation (MV), ECMO treatment, LPS administration + MV, LPS administration + ECMO. As reproducibility is an important criterion for ground-truth labels, we tested the reproducibility of our scoring system by repeating the scoring with observers not involved in the original study (termed validation cohort, “B”). We observed a high similarity of the median scores between the original “A” cohort and the validation “B” cohort (**Figure 1b-d**). We then compared these scores to those assessed with the Matute-Bello system modified for use with digital slides (“C” cohort), normalizing the scores from both scoring systems to a common range. We found that the “C” cohort scores were overall higher and distributed in a bimodal pattern (**Figure 1 b-e)**. To further assess how much the scores from the “B” and the “C” cohort differed from our originally reported scores, we plotted the scores for the “B” and “C” cohorts *versus* the “A” cohort and calculated the slope of the linear regression line and the correlation metric, R-squared (**Figure 1d**). While the “B” cohort showed a high correlation with the “A” cohort (slope of 0.91 and an R-squared value of 0.93), the “C” cohort had a slope of 1.146 which indicates that the Matute-Bello score system skewed the total score towards higher values compared to the two cohorts using the Silva Silva *et al.* system (**Figure 1d**). Overall, our findings indicated that our recently developed semi-quantitative scoring system provides similar results when comparing median values from two cohorts of observers (14). This, together with a large scoring range that allows for nuanced scoring, suggested that it would be well-suited to generate ground-truth labels for supervised training of deep neural networks.

### 3.2 Variability between human scorers reinforces the need for automatization of the scoring with deep neural networks

Based on the notable differences in the distribution of total scores among the two scoring systems, we next sought to assess whether the expertise level affected the sensitivity of either scoring system (**Figure 2**). While there was a generally higher rank correlation between observers with the Silva *et al.* system (**Figure 2c, Supplemental Figure 4a**) as compared to the Matute-Bello system (**Figure 2f**), we detected lower rank correlation based on experience level in both score systems (**Figure 2c, f, and g-i**). Interestingly, we observed higher inter-observer variability between board-certified pathologists with extensive clinical training as compared to the inter-observer variability between observers with a high level of expertise in histological scoring for preclinical models (**Figure 2h)**. Except for one of the board-certified pathologists and one of the novice observers (N1), all other observers had at least 82% rank correlation with one another (**Figure 2c and Supplemental Figure 4a**). In general, the modified Matute-Bello system (modified for digitally scanned slides and scoring at the slide level) resulted in higher inter-observer variability (**Figure 2f, 2i**) and we observed tendencies towards higher scores with a bi-modal distribution between high and low scores for all observers (**Figure 2e**). One limitation of these experiments was that different observers scored the slides using the two different scoring systems, and only a small subset of observers scored using both scoring systems (**Supplemental Figure 1**).

**Figure 2.**
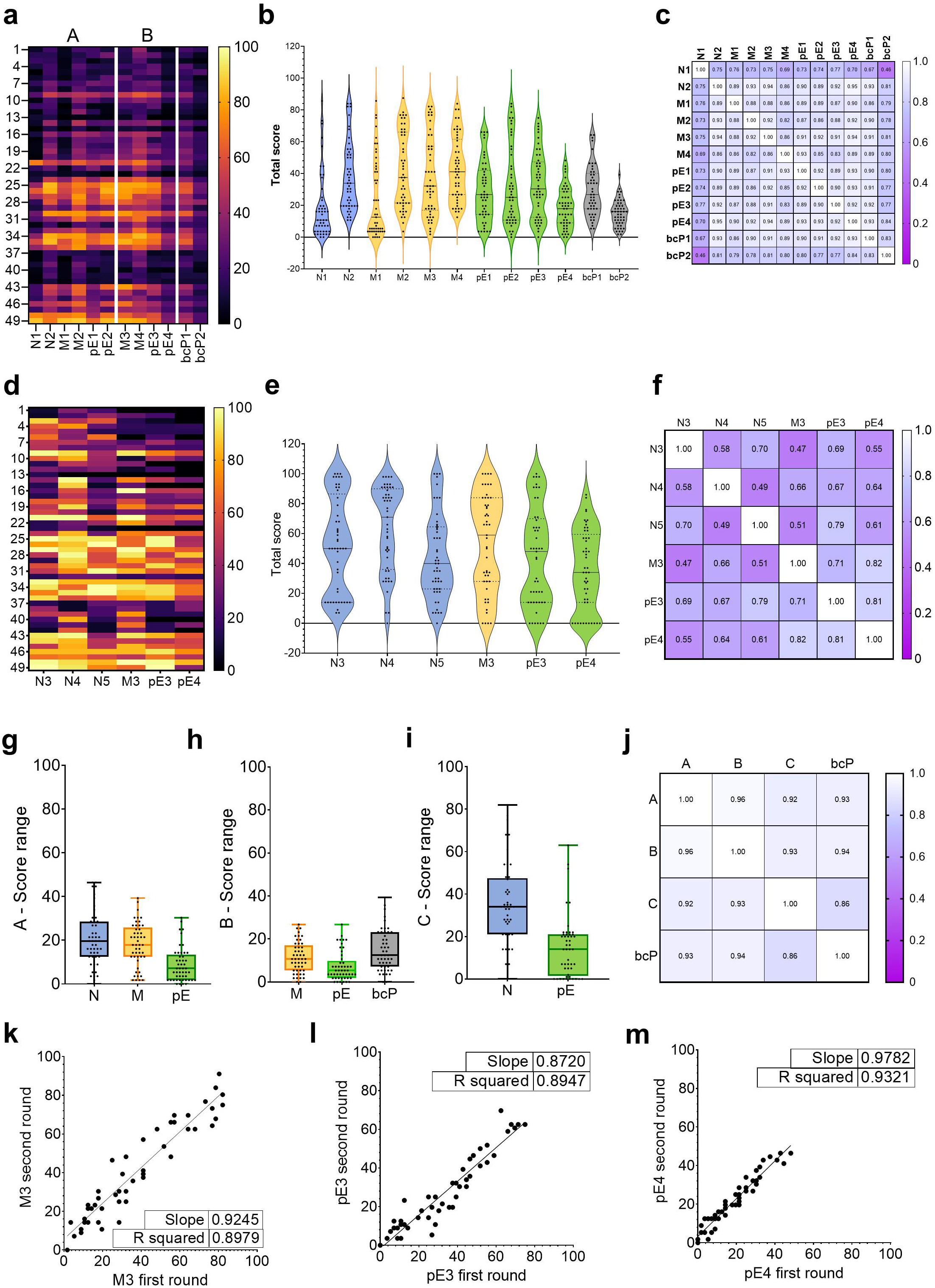
Low agreement between human scorers reinforces the need for automated histological scoring of acute lung injury by deep neural networks. **(A)** Heatmap of normalized total scores from the reviewers in different cohorts that performed scoring based on the Silva *et al.* score system. The lettering on top signifies the original (A) and validation cohort (B). **(B)** Violin plot of the normalized total scores from the reviewers in different cohorts that performed scoring based on the Silva *et al.* score system. **(C)** Correlation matrix showing the Spearman rank correlation between the total scores from the reviewers in different cohorts using the Silva score system. **(D)** Heatmap of normalized total scores from the reviewers that performed scoring based on the Matute-Bello score system. **(E)** Violin plot of the normalized total scores from the reviewers that performed scoring based on the Matute-Bello score system. **(F)** Correlation matrix indicating the Spearmann rank correlation between the total scores from the reviewers that performed scoring based on the Matute-Bello score system. **(G)** Range of the total normalized scores from the Silva score system given by scorers of the original cohort with the same level of expertise. For this and panels H and I, the range was calculated for each slide by subtracting the lowest score given by a reviewer in the respective expertise group from the highest score given by a reviewer in the same expertise group. **(H)** Range of the total normalized scores from the Silva score system given by scorers of the validation cohort with the same level of expertise. **(I)** Range of the total normalized scores from the Matute-Bello score system given by scorers with the same level of expertise. **(J)** Correlation Matrix indicating the correlation between the total normalized median scores from each cohort as well as the group of two certified pathologists the. **(K)** Scatter plot comparing the self-agreement for the total normalized scores from selected reviewer with moderate expertise level (M3), **(L)** pre-clinical expert (pE3) and **(M)** second pre-clinical expert (pE4) for scoring performed on identical slides but over at least one month time period. Slope refers to the slope of the regression lines.

While low inter-observer variability is important to consider when selecting a scoring system for training CNNs, low intra-observer variability is also desired (*i.e.* self-agreement). Therefore, we asked a subset of observers with moderate to high levels of expertise in evaluating pre-clinical models of acute lung injury to repeat their scoring after at least one month using the same set of images. We found that all three observers (**Figure 2k-m**) exhibited high correlation (>87%) between their first and second round of scoring, thus indicating that the Silva *et al.* system has low inter and intra-observer variability making it a promising candidate for training CNNs using a slide-level scoring approach.

### 3.3 Deep neural networks can be trained to score lung histology images

Even though the Silva *et al.* system was reproducible and reliable when comparing the performance of the three cohorts (Cohort A, B, and C), we still observed a relatively high degree of inter and intra-observer variability. Therefore, tools for automated computational scoring would not only cut down scoring time and associated costs but also improve reproducibility and could provide ‘expert’ level scoring to researchers who do not have easy access to observers with a high level of relevant and specific expertise. To explore the feasibility of automated scoring, we trained a series of deep CNNs with a goal of image classification (**Figure 3a-c**). The dataset contained several images for each slide (**Supplemental Figure 2a**), with all images from a slide scored as a group and thus receiving the same label. To fit the required input format of CNNs, images were cut into non-overlapping tiles of 224×224 pixels. Border areas that did not fill an entire tile were discarded (**Supplemental Figure 2b**). All tiles coming from multiple images of the same slide retained the label given to the respective slide which was based on the median score of five reviewers (N1, M1, M2, pE1, and pE2) as ground-truth labels (**Figure 3a**). Such an approach has been successfully used in recent studies using histological images from virtual slides to train deep learning networks for histopathological classification (40). 8073 labelled tiles, between 1113 and 2331 per experimental group, were generated in this manner (**Figure 3c, Supplemental Figure 2a**). After evaluating the distribution of tiles and their corresponding ground truth label, we chose a trimodal classification of the histological images to represent 3 levels of damage: low (non-normalized Silva *et al.* system ≤ 10), medium (10 < non-normalized Silva *et al.* system ≤ 25), and high (non-normalized Silva *et al.* system> 25. These class boundaries were set by considering the distribution of the median scores across the dataset (**Figure 3b)** and the number of tiles (**Figure 3c)** in each experimental group, with the goal of obtaining approximately similar numbers of tiles in each damage class to fit a trimodal distribution. Using these class boundaries, 3368 tiles had a low score, 2668 had a medium score and 2037 had a high score (**Figure 3c**), making the dataset slightly imbalanced.

**Figure 3.**
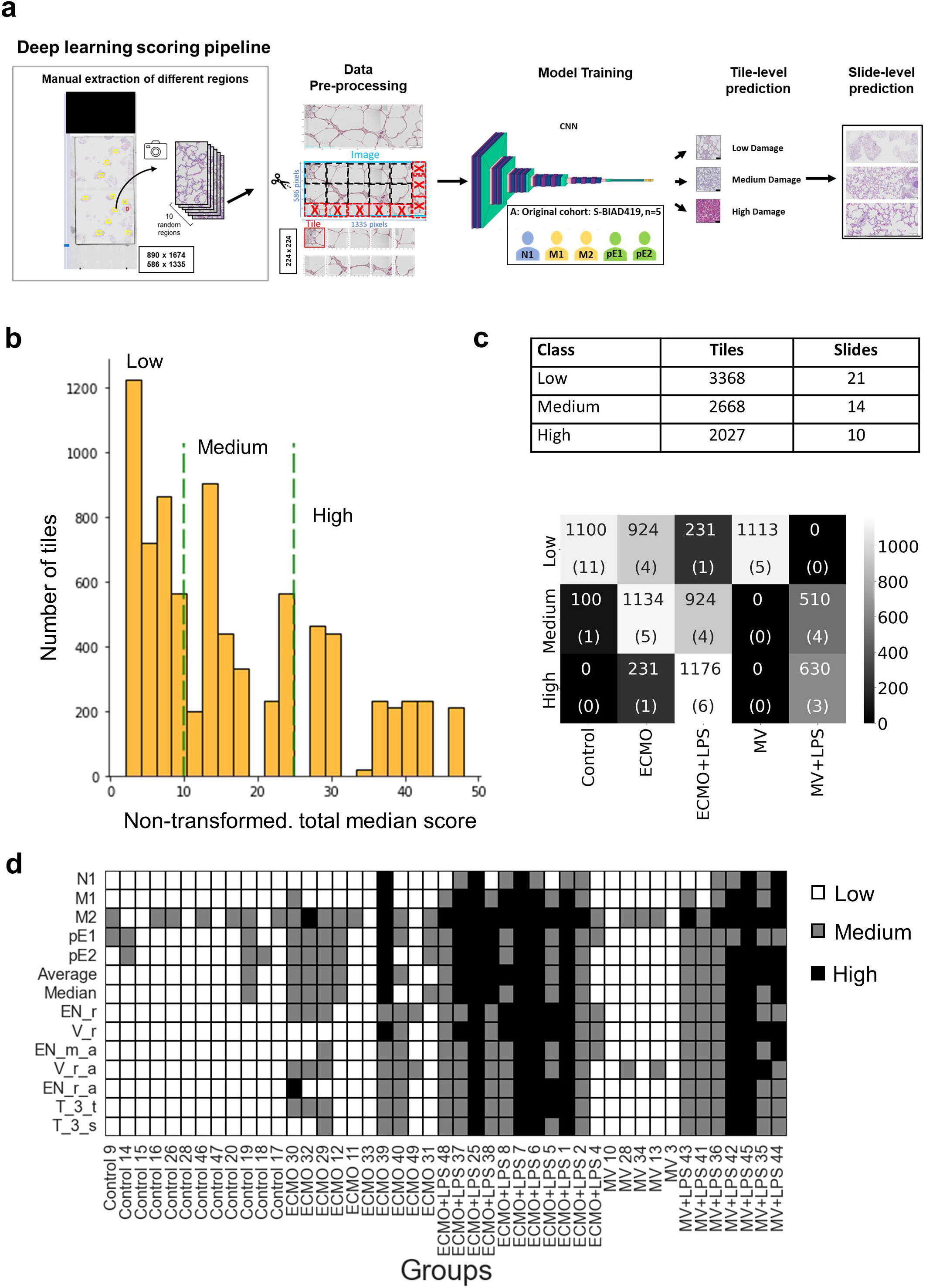
Development of deep neural networks for automated histology scoring. **(A)** Overview over the development of the deep learning scoring pipeline. CNNs were trained on 224 x 224 tiles (with or without augmentation) cut from up to 10 images from each slide with ground truth labels for low, medium and high damage derived from the median slide scores of five reviewers from the original cohort. After training, models were used to make predictions on the tile level which were then aggregated to slide level by “majority vote”. **(B)** Histogram displaying the distribution of tile scores in the dataset used to train the CNN models. Each tile received its score, based on which the ground-truth class label (“low”, “medium” or “high” damage) was later assigned, by calculating the median of the scores given by the five reviewers to the corresponding slide. **(C)** Distribution of scores across classes and treatment groups (MV stands for mechanical ventilation). Median tile scores were converted to 3-class labels, corresponding to low (≤10), medium (10–25) and high damage (>25). **(D)** Slide scores assigned by the reviewers from the original cohort in comparison with predictions by CNNs. Individual reviewer scores, their average and median (used as ground truth label for training the CNNs) and predicted scores by the 2 individual CNN models trained on raw dataset (EN_r and V_r), 3 best individual CNN models trained with augmentation (EN_m_a,V_r_a, and EN_r_a) and two ensemble predictions obtained by combining predictions from the three models on tile level (T_3_t) and slide level (T_3_s). Slides are identified by treatment group and slide number.

Due to the relatively small size of the dataset, splitting into training, validation and test set was not deemed suitable. CNNs (e.g. DenseNet) were initially trained from scratch for binary classification of damage or no damage, but as expected from the small size of the dataset, this led to poor performance (*data not shown*). Instead, 3-fold cross validation was performed for trinary classification (i.e. 3-class). First, the dataset was split into three parts for each CNN type investigated to ensure distribution of experimental groups and damage classification at the slide level. Next, we independently trained three CNNs, each with two of these parts as training set and the third part used as test set. Metrics from each of the three CNNs were then averaged across the three folds. For each fold, all experimental groups and damage classes were present in both the training and test-set and there was no overlap between the test sets (i.e. no overlap between the folds). All tiles derived from the same slide were kept together so that they were either in the training or in the test set to prevent data leakage. As a result, the number of tiles in each fold varied slightly (**Supplemental Figure 2c**).

Due to the small number of images available for training, we opted for transfer learning, which means that weights pretrained on ImageNet were used for all layers except the top layers and then kept frozen while the top layers were fine-tuned on the non-augmented dataset (these CNNs are referred to as “raw data” models hereafter). Two types of pre-trained CNN architectures were selected for testing: VGG16 and EfficientNetB4 (**Supplemental Figure 3b**), due to the fact that they are both large networks which are widely employed for image classification and have shown good performance across diverse image classification tasks (37, 41). After-fine tuning, raw data VGG16 (model V_r) and EfficientNetB4 (model EN_r) models were quite similar in their performance with average weighted F1 scores of 0.75 and 0.73 across the three folds, respectively (**Figure 3d, 4a, Supplemental Table 3**). All misclassified slides occurred between the medium and the high or low class, whereas no misclassifications were seen between the high and the low class (**Figure 4a**).

**Figure 4.**
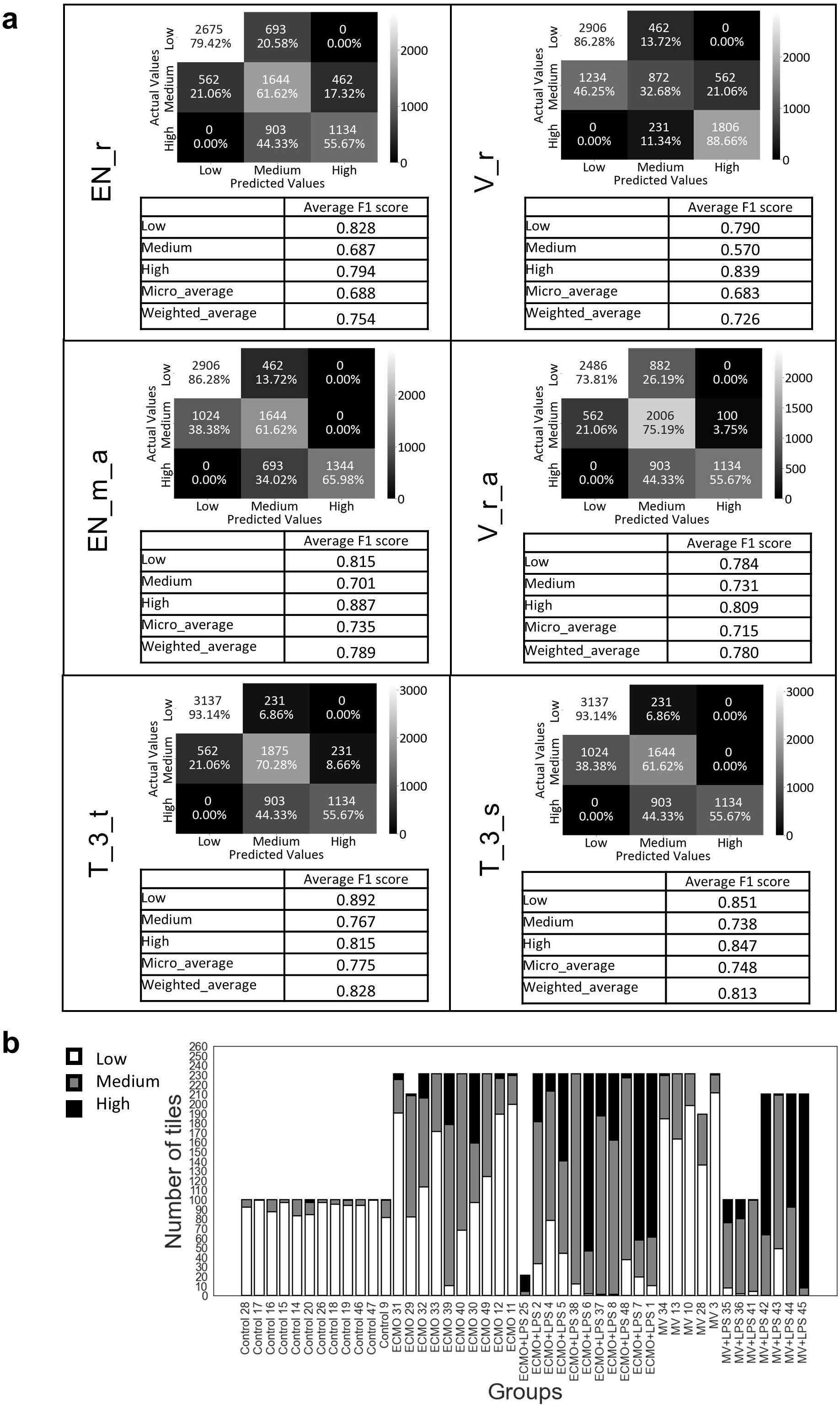
Evaluation of the trained CNNs. **(A)** Summary of CNN evaluations. Results obtained from combining the evaluation for all three folds are shown for Efficient (indicated with EN_ prefixes), and VGG16 (“V_”) CNNs trained on the raw dataset (EN_r and V_r), augmented dataset (two bests are shown, EN_m_a and V_r_a) and two ensemble models. For the ensemble models a majority vote on tile level (T_3_t) or slide level (T_3_s) was used to aggregate the slide label prediction of the three best models. The confusion matrices show true tile labels vs predicted tile labels for all damage classes. The total number of tiles calculated by adding the corresponding matrices from the three folds is shown in the top of each cell and the cell shaded correspondingly as indicated by the color bar. The number of tiles normalized as percentage of the true labels is shown in the bottom of each cell. Values were calculated with the “confusion_matrix” function from scikit-learn (v. 1.0.1). The tables underneath the matrices show the averages of the F1 score for each class, of the micro averaged F1 score and of the weighted average F1 score. Note that the values in the tables are not calculated form the confusion matrices but instead by averaging the respective slide-level metrics from the three folds. **(B)** Tile scores predicted by the best individual model (EN_m_a) for each slide. Each bar represents one slide with multiple images from which the tiles were cut. The height of each bar represents the number of tiles that received the respective score. Bar heights vary because the number of images and their dimensions differed between slides resulting in a different overall number of tiles.

### 3.4 Dataset augmentation and model ensembles improve scoring performance

When training with small datasets such as the one available for this study, augmentation can improve performance. We therefore evaluated different strategies to augment the dataset (**Supplemental Table 1**). Blurring and brightness variation were explored due to the fact that they occur commonly in histology data and can impact the performance of deep learning networks (42). Further, we also used rotations, shifting and flipping. To determine a suitable blurring kernel, different sizes were tested (5 x 5, 10 x 10, 25 x 25 and 40 x40) (**Supplemental Figure 3a** and *data not shown*). The larger kernel sizes were deemed to blur the images to levels not typically observed in histology data and thus 5 x 5 kernels were chosen for augmentation. The introduction of blurring decreased the average weighted F1 scores of both the VGG16 (model V_b) and EfficientNetB4 (model EN_b) model (**Supplemental Table 3**) In contrast, a combination of change of brightness, flipping, shifting and slight random rotation (from -20 to 20 degrees) improved the average weighted F1 score of the EfficientNetB4 model to 0.79 and a combination of change of brightness, flipping and random rotation (from -90 to 90 degrees) improved the average weighted F1 score of the VGG16 model to 0.78 (**Figure 3d and 4a, Supplemental Table 3**). In both cases this was due to an improvement in both precision and recall. As with the previous models, no misclassifications between the high and low classes were observed (**Figure 4a**). For both models trained with the raw dataset and models trained with augmentation, relatively large differences were seen between the results for the three different folds (**Supplemental Table 3**).

Another strategy to improve the performance of deep neural networks is to aggregate the predictions of several models in an ensemble. To evaluate the benefit of an ensemble strategy, we first aggregated the slide-level predictions of the three best models (EN_m_a, EN_r_a, VGG16_r_a) using a majority vote approach, with a medium score assigned in case all models scored differently. The slide-level ensemble (T_3_s) had a weighted average F1 score of 0.81, which was higher than the scores of the individual models (**Figure 3d, 4a**). The ensemble approach also reduced the number of misclassified slides to 9 whereas the individual models had 10 or 12 misclassified slides. Notably, all three models agreed for 7 of the 9 misclassified slides. We next tested an ensemble approach with aggregation on the tile level (model T_3_t) which resulted in an F1 score of 0.83, also with 9 misclassified slides (but only some the same as for the slide-based ensemble).

### 3.5 Deep neural networks can highlight intra-sample heterogeneity and detect subtle damage

Human scoring is typically performed on slide or image level due to time constraints. However, this approach hides the large degree of heterogeneity across a tissue section. In contrast to human reviewers, our CNNs made predictions on the much smaller tiles, which were only later aggregated to slide level scores. Scoring with CNNs therefore makes it possible to examine scores on the tile level as well.

When investigating the tile scores of the best individual model (EN_m_a), we observed that most slides had at least some tiles scored as a class not chosen by the majority vote (**Figure 4b**). When overlaying the tile scores over the full image, we observed that in many cases, the model appeared to correctly identify areas with different levels of damage (**Supplemental Figure 5**). However, it was also clear that upon subjective assessment of predicted tile-level scores and consultation among experts (pE4 and bcP2), some tiles were misclassified as having high damage but instead contained moderate levels of damage at the tile-level (**Supplemental Figure** 5). Nonetheless, the improvement of the models towards correct damage classification was evident in the tile-level visualizations in the models which used augmentation as well as the ensemble approach.

Interestingly, we observed that most of the CNN models identified slides coming from animals treated with ECMO as having a moderate degree of injury as compared to baseline (control) or animals treated with mechanical ventilation (MV). In our original description of the scoring system, total injury assessment by five observers failed to detect differences between ECMO and MV treated animals (13). However, after re-analyzing this data (**Figure 3d**) in more detail, it became apparent that the pre-clinical experts (pE) and one observer with intermediate experience (M2) did indeed identify a higher degree of injury in these slides as compared to baseline and MV. The aggregation of 5 scorers with varying degrees of expertise thus seems to have limited the sensitivity in our original analysis. Differences would have been detected if only the pE scores had been considered. However, it is standard to use multiple scorers in semi-quantitative histology, including those with different levels of histological experience. Thus, it appears that several of the CNNs were able to identify the level of injury to a similar extent as the pre-clinical experts.

### 3.6 Deep neural networks can make valid predictions on an unseen data set

Neural networks can struggle with generalization, that is, they often cannot make valid predictions on images from datasets that have not been part of the training data. To test whether our neural networks could generalize to new datasets, we extracted pig lung histology images from a recent publication which induced lung injuries in both a similar and different manner to the one utilized in S-BIAD419. Lung injury in S-BIAD419 was induced used intravenous and intratracheal administration of LPS derived from *E. coli* while Ref (43) used intravenous administration of live *E. coli* or direct lung injury using volutrauma, hyperoxia and bronchoscopically-delivered gastric particles; a third group combining both injury models was also used in Ref (43). Published images were then processed into tiles in the same way as for our dataset, regardless of the magnification, and predictions made with our best individual neural network (EN_m_a) and the best ensemble model (T_3_t). When visually inspecting overlays of the predictions over the images (pE4 and bcP2), we could see that the neural network correctly identified tiles with low, medium and high tissue damage in many cases (**Supplemental Figure 6**). This suggests that the neural network might be able to generalize relatively well, but future studies with unrelated datasets with ground-truth labels will be needed to quantify this ability to generalize.

## 4. Discussion

This study explored the possibility of using a semi-quantitative score system at the slide-level to train deep neural networks with the goal to automate this difficult, costly and time-consuming human task and increase reproducibility. Originally, scoring systems were developed from photomicrographs, taken from randomly chosen fields with blind microscope stage controls and captured without adjustments (44). During the last few years, sample processing has gone through dramatic modernizations which has standardized many steps to reduce human variability and the amount of time associated with histological processing. This includes the automatization of tissue processing itself, the use of automatic slide staining and digital slide scanning. However, automatization of image analysis is still lacking for many tissues and diseases. Such tools would ideally be used as a standardized tool shared across groups to facilitate the collaboration and comparison across different laboratories. Automation of image analysis would also increase the throughput, and reduce cost, batch and inter-laboratory differences (45).

Although widely used scoring methods exist for analyzing the extent of lung injury in histology samples from preclinical animal models for fibrosis (44, 46) and acute lung injury (9, 10), these methods are challenging to implement for large animal models of ARDS as there are major species specific differences (*e.g.* diameter of the alveoli). As for many other diseases, murine animal models have been widely explored for their versatility, reproducibility and the ease in standardizing and manipulating their genetic background (5, 9, 13). Furthermore, their lower cost allows for the study of multiple variables potentially related to disease (*e.g.* effects of age, co-morbidities and biological sex) (47–49). However, for animal-based studies that aim to evaluate disease progression and/or potential therapies, incorporation of clinical standards of care, or potential therapeutic interventions such as mechanical ventilation or ECMO, large animal models (typically porcine or ovine) are the most appropriate because their lung anatomy and injury features are more similar to humans and incorporation of surgical interventions are possible (50, 51). Additionally, the increased cost associated with their use and technical challenges in implementing these model are associated with the fact that fewer experiments have been performed and thus there is less knowledge around the role of some experimental variables, such as biological sex (43), which have been shown to be important clinically (52). However, in comparison to small animal models, large animal models offer the potential for serial biopsies which can allow for comparison to clinical variables. This is particularly important as the main histological pattern of ARDS, termed diffuse alveolar damage (DAD), is not observed in all patients with clinically confirmed ARDS (53). Therefore, large animal models provide a unique opportunity to gain insight into the histological progression of the disease, which is technically challenging to obtain in the clinical setting (54). It is consequently of importance to have semi-quantitative scoring systems, ideally automatized, that work reliably in such large animal models (50, 55).

Existing semi-quantitative scoring systems typically use a set of injury features for which the extent of occurrence is judged by several human scorers. There has been variation not only in the injury features that are included but also in the numerical range used to score the injury. Ranges such as 0-2, 0-3, 0-5 and up to 0-8 have been employed (44, 46, 56–61). A smaller range may compromise the sensitivity of the score, as it masks the heterogenicity of the samples and does not allow for nuanced scoring, while too large ranges might affect reproducibility, as the distinction between different degrees of injury becomes less obvious (44, 46, 62). When using human scoring to train deep neural networks, it is essential that the semi-quantitative system used by the human scorers to generate ground truth labels is reliable. We therefore explored the reproducibility of our recently developed Silva *et al.* system (13, 14), comparing different cohorts of scorers as well as scorers within our validation cohort. The variability between individual scorers was relatively large, even between experts, underlining the need for automatization of this essential analysis step. However, when grouping several scorers into a cohort we found high reproducibility between our original and validation cohorts, which comprised different persons. This indicates that our score system was reproducible and reliable enough for training deep neural networks.

As a proof-of-concept, we demonstrated that categorical labels derived from the median total injury score of five human scorers were indeed suitable for training CNNs that can distinguish between low, medium and high tissue damage to a degree comparable with human scorers. One limitation was the relatively small size of the dataset, which necessitated the use of a cross-fold validation strategy (63). Relatively large differences were seen between the results for the three different folds. This was likely due to differences in the composition of the folds, and because with small datasets, the influence of individual slides is relatively large. One aspect to consider when judging the performance of the models is that despite the use of five scorers, the ground truth annotations might have been incorrect at least in some cases due to the difficulty of the scoring task and the high variability seen between scorers. Interestingly, the agreement between different models was higher than the agreement between human scorers and notably all three of the best models agreed for 7 of the 9 slides misclassified by the slide-level ensemble. This could suggest that the variations between the model predictions were more similar than the variation across the different observers scorers.

The damage observed across a tissue section is very heterogenous with many features being very unevenly distributed and thus hard to judge. Because human scorers did not score on tile level due to the enormous amount of time required for this task, all tiles inherited their “ground truth” label from scores assigned at the slide level as has been done in previous studies utilizing deep learning for histopathological classification from virtual slides (40). This means that a significant portion of slides had “ground truth” labels that may not have been correct. In addition, the use of a trinary classification system and the class boundaries we selected may not have been ideal. Nonetheless, qualitative examination suggested that the models often correctly identified areas of the images that showed different degrees of damage (i.e. low, medium or high damage), but there were also some tiles which qualitatively appeared to be mis-classified when reviewed and discussed by experts (pE4 and bcP2) (**Supplemental Figure 5**). Further work is needed in order to improve the accuracy at the tile-level, which may also require optimizing how the damage classes are assigned and which labels are used as ground truths. As it is difficult for humans to score damage that is heterogenous across a large image area, it is also possible that at least some of the “errors” made by the models were actually correct (64). To explore this further, a larger group of expert level scorers would need to score more slides and scoring should be performed on smaller image sections as has been previously described to be ideal for preclinical animal models (9).

The fact that models can generate more fine-grained scoring on tile-level is another large advantage of their use over human scorers. Such fine-grained scoring is a time-intensive task for humans which not only makes it costly and often not feasible but also increases the possibility of error due to fatigue. Tile-based scoring by CNNs would make it possible to investigate differences in the spatial distribution of damage, for example, over the course of the disease, or across different forms of ARDS and individuals, which is currently poorly understood (65, 66). Tile-based scoring might potentially also be able to reveal more subtle differences between experimental groups. Such subtle differences could for example result from mild pre-existing lung injury, age or sex-specific differences which could result in different inflammatory responses. The interplay of these individual differences are currently poorly understood, especially in large animal models and at the molecular level (48, 67–69). Due to the small dataset size in this proof-of-concept study, it was not possible to investigate whether the CNN models had any biases related to this, but future studies should aim to compile training datasets are balanced and large enough to investigate these important issues.

Histology scoring is a bottleneck in many research groups. Using an automated neural network-based scoring system to complement human based histologic assessment could help accelerate research and bring down costs. At the same time, the large variations in scoring between experiments, individuals and research groups, and even between repeated scoring rounds by the same people, which are widely recognized to constitute a problem, could be largely eliminated with a community-adopted automated scoring tool (63). However, scoring systems based on CNNs or CNN ensembles, which performed even better, not only have the potential to outperform human scorers in terms of cost, speed and reproducibility but they are also better suited to detect subtle differences and intra-sample heterogeneity. Tile-based scoring with CNNs makes it possible to gain more nuanced insights into lung injury by exploring spatial variability which simply remains hidden in most studies using human scorers due to the enormous amount of labor this would require.

Despite the clear advantages of CNN-based scoring and the successful proof-of-concept in this study, challenges in training these networks remain, such as the limited size of available annotated data, potential biases in existing datasets, unreliable ground-truth and a lack of fine-grained tile-based annotations. Since the performance of CNN-based classification models is strongly affected by these limitations, we expect that our models can be improved even further in the future by investing in the production of high-quality annotated datasets.

## Acknowledgements

We thank all members of the “Cell Death, Lysosomes and Artificial Intelligence” and “Lung Bioengineering and Regeneration” groups for helpful comments throughout the development of this project as well as all of the persons who served as observers. We are especially grateful for Dennis Medved (Lund University) who co-supervised the undergraduate students August Lidfeldt and Ludwig Hedlund. This project is partially funded by Swedish Research Council Starting Grants (D.E.W. Dnr 2018-02352, S. A. Dnr 2016-02003). The Knut and Alice Wallenberg foundation is acknowledged for generous support (D.E.W). This study was also supported by grants to Science for Life Laboratory from the Knut and Alice Wallenberg Foundation (S.A. 2020.0182), which were distributed through the SciLifeLab and KAW National COVID-19 Research Program. The Swedish Foundation of Strategic Research is acknowledge for their support through a research grant awarded to V.S. (UKR22-0081).

Computations and data handling was enabled by resources provided by the Swedish National Infrastructure for Computing (SNIC), partially funded by the Swedish Research Council through grant agreement no. 2018-05973, specifically the Berzelius resource provided by the Knut and Alice Wallenberg Foundation at the National Supercomputer Centre at Linköping University, the High Performance Computing Center North (HPC2N) at Umeå University, LUNARC at Lund University, and Alvis at the Chalmers Centre for Computational Science and Engineering (C3SE). Pilot experiments were conducted with Google Colaboratory, provided by Google Research.

## CRediT author statement

Iran A. N. Silva: Conceptualization, Methodology, Data Curation, Formal analysis, Investigation, Writing original draft, Visualization

Salma Kazemi Rashed: Conceptualization, Methodology, Software, Formal analysis, Investigation, Writing - Original Draft, Visualization

Nika Gvazava: Data Curation, Formal analysis, Validation, Investigation, Writing - Review & Editing

Valeriia Skoryk: Formal analysis, Investigation, Validation, Writing - Review & Editing

John Stegmayr: Methodology, Formal analysis, Writing - Review & Editing

August Lidfeldt: Formal analysis, Software, Investigation, Writing - Review & Editing

Ludwig Hedlund: Formal analysis, Software, Investigation, Writing - Review & Editing

Sonja Aits: Conceptualization, Methodology, Resources, Writing - Original draft, Writing - Review & Editing, Visualization, Supervision, Project administration, Funding acquisition

Darcy Wagner: Conceptualization, Methodology, Formal analysis, Validation, Investigation, Writing - Original draft, Visualization, Supervision, Resources, Project administration, Funding acquisition

## Notes

https://www.ebi.ac.uk/biostudies/bioimages/studies/S-BIAD419

### Competing Interest Statement

The authors have declared no competing interest.

