## Supplemental for "Deep learning for rapid and reproducible histology scoring of lung injury in a porcine model"

**Supplemental Figure 1. Overview of experimental approach for scoring of slides in S-BIAD419 with observers of various expertise and their correlation with one another.** Two different scoring approaches were used (Silva and Matute-Bello as described in Methods). The heading “A” represents scores deposited in S-BIAD419 as metadata while the subsequent three columns represents observers which scored slides in the current manuscript. “B” = observers who performed similar scoring as done in “A". "Repeat" = group of reviewers from "B" who repeated the same manual scoring task more than one month later. “C” = observers who scored using Matute-Bello on the same set of slides. ID represents individual scorers. The first letter designates experience level as detailed in Figure 1A while the number is the reviewer number within that experience category.

| ID | Expertise Level | A | B | Repeat | C |
|---|---|---|---|---|---|
| **N1** | Novice | x | | | |
| **N2** | Novice | x | | | |
| **M1** | Moderate | x | | | |
| **M2** | Moderate | x | | | |
| **pE1** | Expert | x | | | |
| **pE2** | Expert | x | | | |
| **N3** | Novice | | | | x |
| **N4** | Novice | | | | x |
| **N5** | Novice | | | | x |
| **M3** | Moderate | | x | x | x |
| **pE3** | Expert | | x | x | x |
| **pE4** | Expert | | x | x | x |
| **M4** | Moderate | | x | | |
| **bcP1** | Board certified Pathologist | | x | | |
| **bcP2** | Board certified Pathologist | | x | | |

**Supplemental Figure 2. Overview of the dataset used for the CNN training.**
**a)** Characteristics of the dataset extracted from S-BIAD419 used for CNN training. **b)** Data pre-processing for CNN training. The images were cut into adjacent, non-overlapping tiles of pixel size 224 x 224 to match the input size expected by the CNNs. Tiles not fully covered by images were discarded. This also removed areas of the larger images that contained text or size bars. **c)** Number of tiles in the test and training sets per fold and treatment group

**a**

| Treatment group | Image dimensions | Number of slides | Number of images | Number of generated tiles |
|---|---|---|---|---|
| Control | 586 x 1335 (RGB) | 12 | 120 | 1200 |
| ECMO | 817 x 1635 (RGB) | 10 | 109 | 2289 |
| ECMO+LPS | 817 x 1635 (RGB) | 11 | 111 | 2331 |
| MV | 817 x 1635 (RGB) | 5 | 53 | 1113 |
| MV+LPS | 586 x 1335 (RGB) | 7 | 30 | 300 |
| | 890 x 1674 (RGB) | | 40 | 840 |

**b**

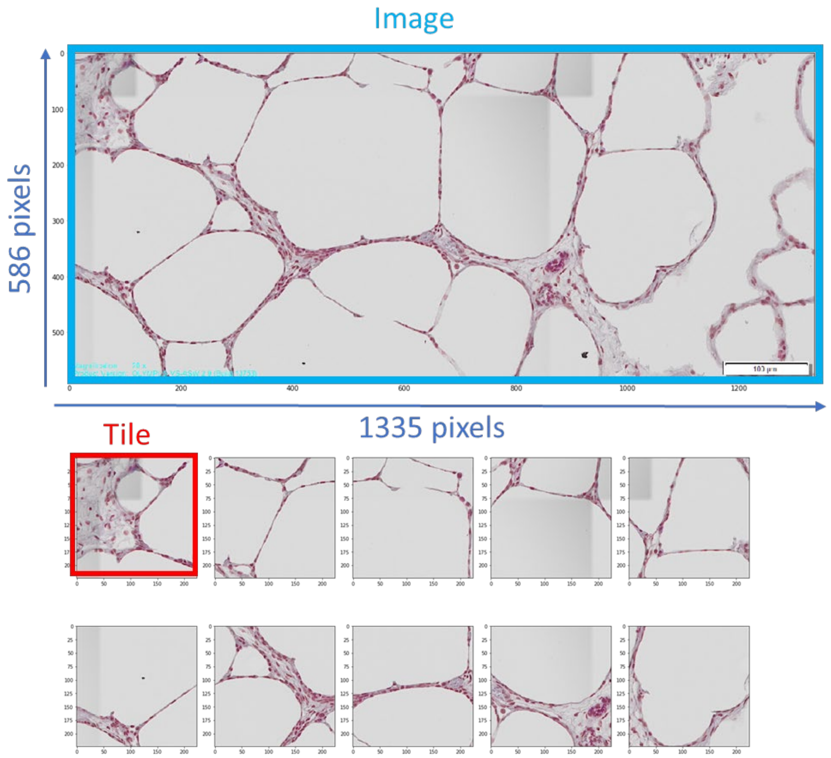


**c**

| | | fold1 | fold2 | fold3 | sum |
|---|---|---|---|---|---|
| Test | Low | 1093 | 762 | 1513 | 3368 |
| | Medium | 872 | 1355 | 441 | 2668 |
| | High | 231 | 672 | 1134 | 2037 |
| Train | Low | 2275 | 2606 | 1855 | 6736 |
| | Medium | 1796 | 1313 | 2227 | 5336 |
| | High | 1806 | 1365 | 903 | 4074 |

**Supplemental Figure 3. Overview of the deep learning approach. a)** Examples of augmentations used to increase the size and variation of the CNN training data. **b)** VGG16 with top layer replaced by the following architecture: Fully Connected (Dense) (128) + Dropout (0.5) + Dense (SoftMax) layers with 3 classes at last layer. Only these new layers were fine-tuned whereas the other VGG16 layers. with weights from pretraining on ImageNet. remained frozen (approximately 3000000 trainable parameters). The network schematic is plotted with the visualkeras (0.0.2) python package. **c)** EfficientNetB4 with top layer replaced as for the VGG16. Only the new layers were fine-tuned whereas the other EfficientNetB4 layers with ImageNet weights remained frozen (approximately 11000000 trainable parameters).

a

b

c

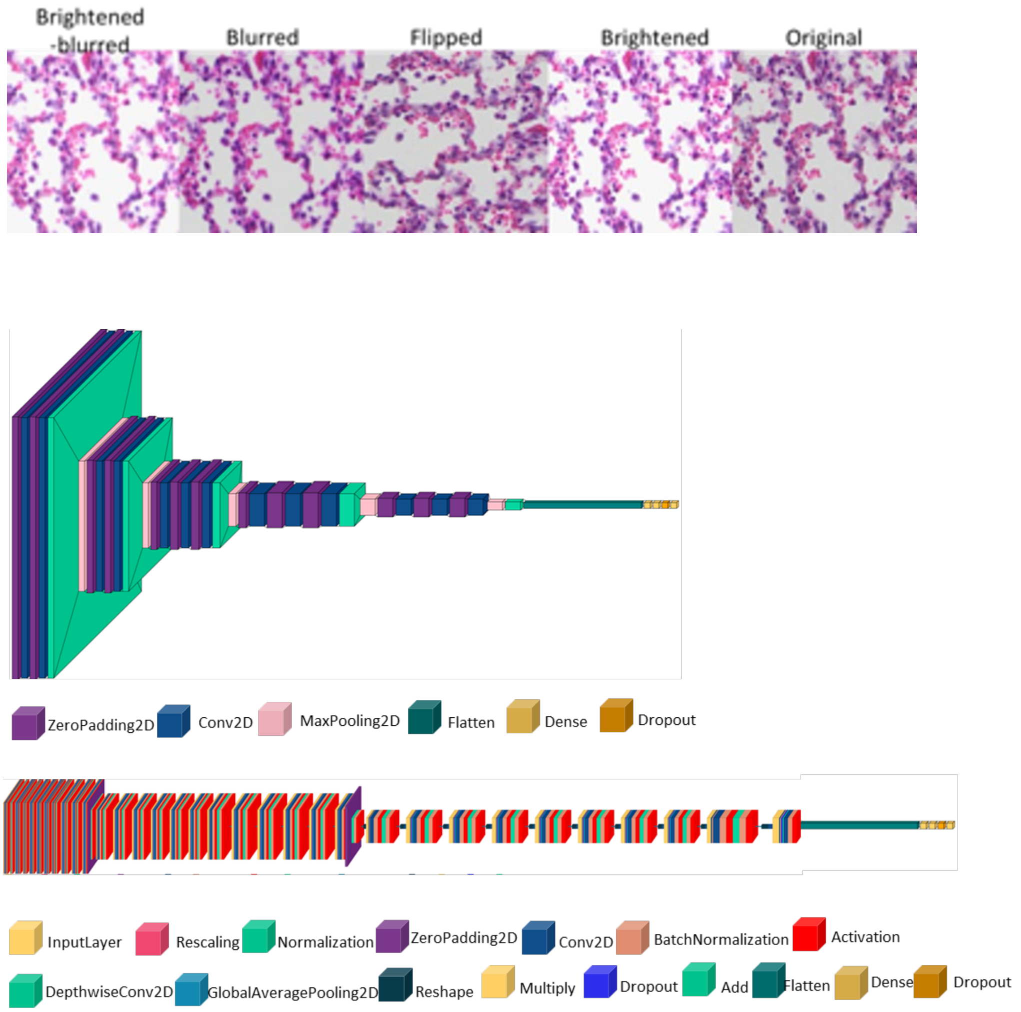

**Supplemental Figure 4. Spearman's rank correlation matrix among different observers across cohort's A and B and compared to board certified pathologists (bcP) and among the different scoring cohorts. a)** Labels for rows and columns correspond to experience level (prefix) and observer identity (number) as outline in Figure 1A and Supplemental 1A. This is an enlargement of Figure 2C in the main images. **b)** Spearman's correlation matrix comparing the median value of each slide with the value given in the additional scoring cohort. Cohort A (S-BIAD419) and cohort B (this paper) scored with the score system described by Silva et al. *Bio-protocols* 2022 while cohort C used a modified version of Matute-Bello et al. *AJRCMB* 2011; other definitions: bcP= board certified clinical pathologists. Repetition = three observers in cohort B scored the same slides at least one month apart

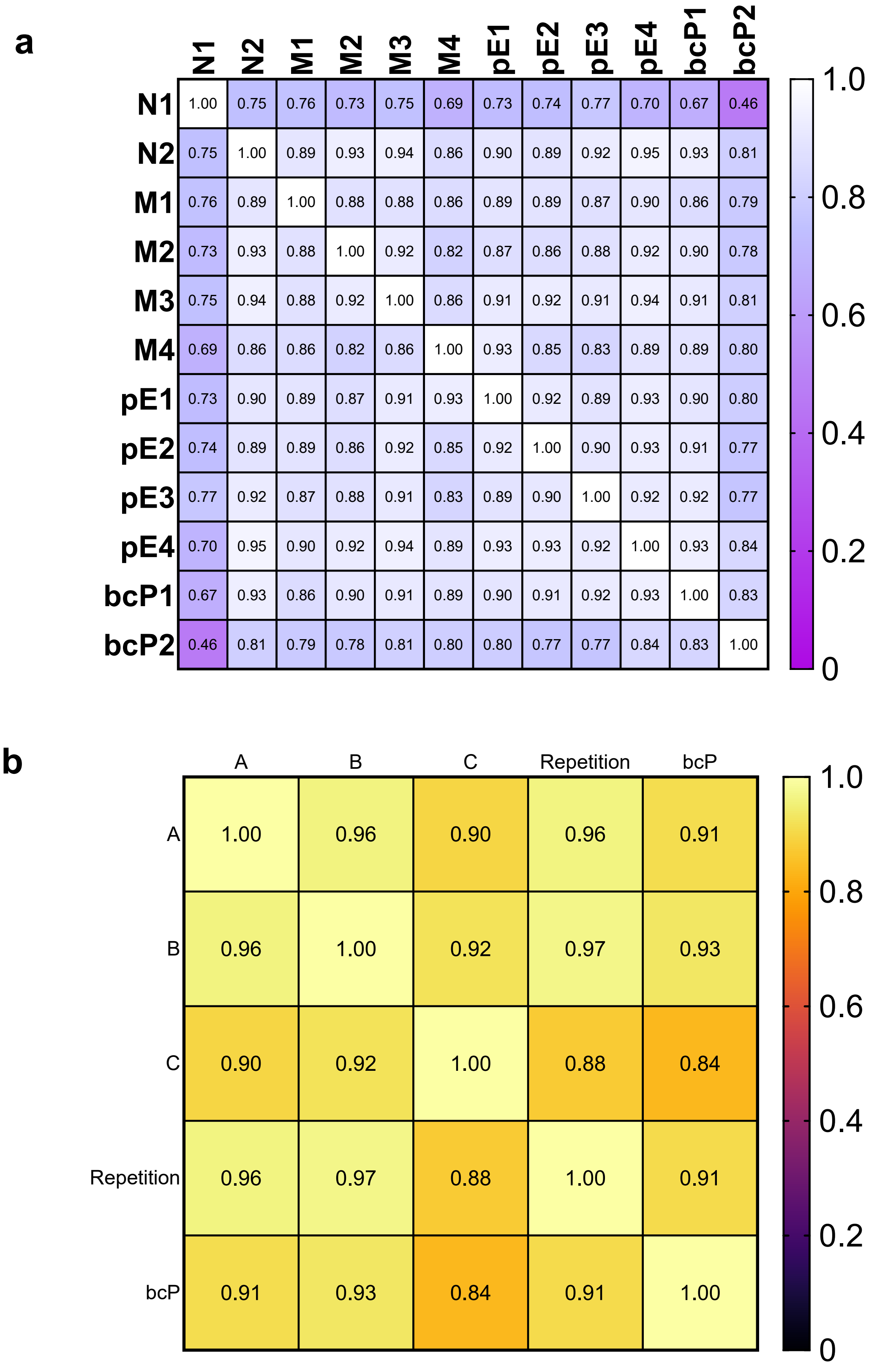

**Supplemental Figure 5. Tile level predictions by CNNs highlight heterogeneous damage across histology images.** All extracted images and tiles inherited the median total score from human based scoring at the slide-level (top row). Based on the total score for each slide, all tiles in that extracted image were assigned a gold-standard ground truth label (low, medium, or high damage based on the thresholds set in Figure 3b). CNN models were then used to predict the scores at the tile level (below each model). Tile scores from the three best models were then aggregated into an ensemble model (T_3_t) using the majority vote strategy. Tile scores predicted by the ensemble model are indicated by tile shading. with light grey for low damage, medium grey for medium damage, and dark grey for high damage. Predicted slide level damage from the tile-level predictions are shown in bold by majority vote.

| Model | Medium Damage<br>Human Slide-Level Score: 21/56 | High Damage<br>Human Slide-Level Score: 39/56 |
|---|---|---|
| **En_r** | 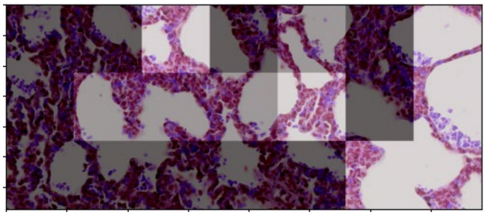 | 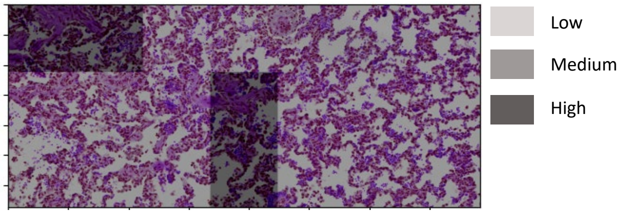 |
| **Majority Vote** (image-level) | **High Damage** (11/21 tiles)<br>Medium Damage (4/21 tiles)<br>Low Damage (6/21 tiles) | High Damage (4/21 tiles)<br>**Medium Damage** (17/21 tiles)<br>Low Damage (0 tiles) |
| **En_m_a** | 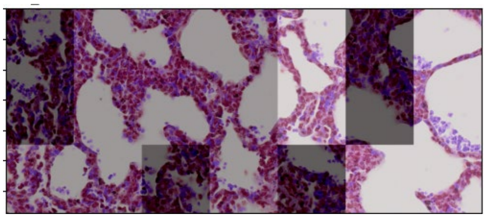 | 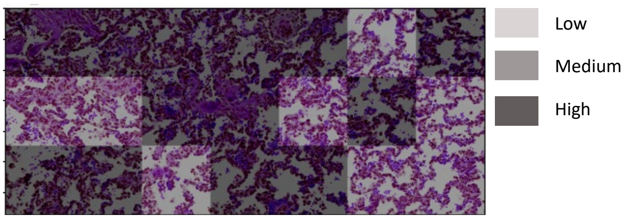 |
| **Majority Vote** (image-level) | High Damage (6/21 tiles)<br>**Medium Damage** (9/21 tiles)<br>Low Damage (6/21 tiles) | **High Damage** (13/21 tiles)<br>Medium Damage (8/21 tiles)<br>Low Damage (0/21 tiles) |
| **T_3_t** | 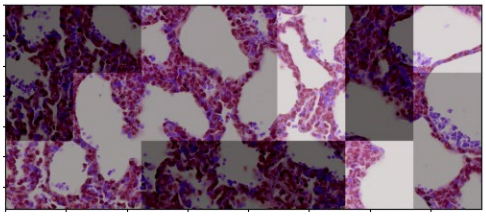 | 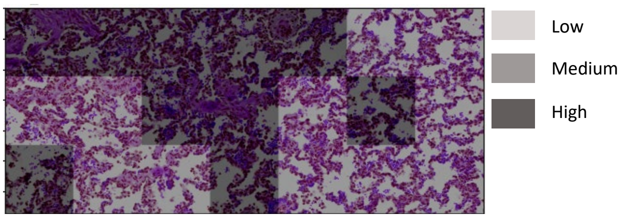 |
| **Majority Vote** (image-level) | High Damage (8/21 tiles)<br>**Medium Damage** (9/21 tiles)<br>Low Damage (4/21 tiles) | High Damage (10/21 tiles)<br>**Medium Damage** (11/21 tiles)<br>Low Damage (0/21 tiles) |

**Supplemental Figure 6 Deep neural networks can make valid predictions on an unseen dataset**. Pig lung histology images (PMID: 33991456) were processed into tiles in the same way as for our dataset, regardless of the magnification, and predictions made with our best individual neural network (EN_m_a) and the best ensemble model (T_3_t) using the majority vote strategy are shown here. Tile scores predicted by the (EN_m_a) and the ensemble model (T_3_t) are indicated by tile shading. with light grey for low damage, medium grey for medium damage, and dark grey for high damage.

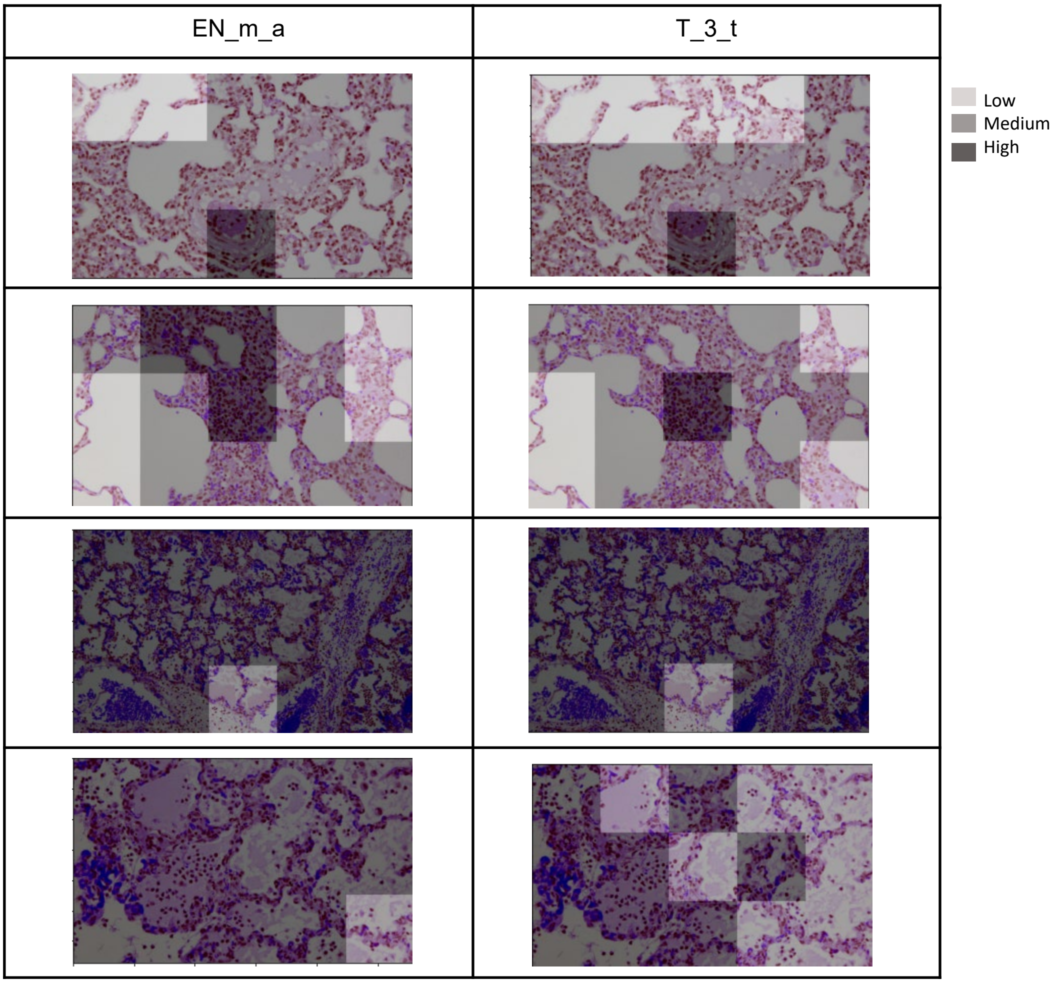

**Supplemental Table 1. Abbreviation of CNN model names and details of the augmentation.** CNN models were trained without ("raw data") or with one of three augmentation forms: A) brightness change (by multiplying pixel values with a random value in range of 0.8 to 1.2) + blurring with 5x5 kernel + horizontal flip. B) brightness change (range 0.8 to 1.3) + rotation in the range of -20 to 20 degrees + flipping + shifting or C) brightness change (range of 0.8 to 1.3) + rotation of 90, 180 or 270 degrees + flipping.

| Abbreviation | CNN model + training data |
|---|---|
| EN_r | EfficientNetB4, raw data |
| V_r | VGG16, raw data |
| EN_b | EfficientNetB4, blurring/brightness change (0.8 to1.2) |
| V_b | VGG16, blurring/brightness change (0.8 to 1.2) |
| EN_r_a | EfficientNetB4, rotation (-20 to 20)/flipping/shifting/brightness change (0.8 to 1.3) |
| V_r_a | Vgg16, rotation (-20 to 20)/flipping/shifting/brightness change (0.8 to 1.3) |
| EN_m_a | EfficientNetB4, rotation (90/180/270)/flipping/brightness change (0.8 to 1.3) |
| V_m_a | Vgg16, rotation (90/180/270)/flipping/brightness change (0.8 to 1.3) |

**Supplemental Table 2. Abbreviation of ensemble CNN models. aggregation rule and the level of decision (tile or slide).** The results of three best models as EN_m_a, EN_r_a, and V_r_a were ensembled by majority rule in tile and slide level score.

| Abbreviation | Ensemble description |
|---|---|
| T_3_t | Top3 best models. majority vote. tile level |
| T_3_s | Top3_best_models. majority vote. slide level |

### Supplemental Table 3. Performance of different CNN's with and without different augmentations.

| | | fold1 | | | fold2 | | | fold3 | | | Average | | |
|---|---|---|---|---|---|---|---|---|---|---|---|---|---|
| | | Precision | Recall | F1-score | Precision | Recall | F1-score | Precision | Recall | F1-score | Precision | Recall | F1-score |
| EN_r | Low | 0.789 | 0.789 | 0.789 | 0.697 | 1 | 0.822 | 1 | 0.695 | 0.82 | 0.829 | 0.828 | 0.828 |
| | Medium | 0.735 | 0.735 | 0.735 | 1 | 0.415 | 0.586 | 0.244 | 1 | 0.393 | 0.66 | 0.717 | 0.687 |
| | High | 1 | 1 | 1 | 0.593 | 1 | 0.744 | 1 | 0.204 | 0.338 | 0.864 | 0.735 | 0.794 |
| | Micro_average | 0.79 | 0.79 | 0.79 | 0.716 | 0.716 | 0.716 | 0.558 | 0.558 | 0.558 | 0.688 | 0.688 | 0.688 |
| | weighted_average | 0.79 | 0.79 | 0.79 | 0.819 | 0.716 | 0.764 | 0.892 | 0.558 | 0.687 | 0.834 | 0.688 | 0.754 |
| EN_b | Low | 0.713 | 1 | 0.832 | 0.576 | 1 | 0.731 | 1 | 0.847 | 0.917 | 0.763 | 0.949 | 0.846 |
| | Medium | 1 | 0.494 | 0.662 | 0.612 | 0.244 | 0.349 | 0.389 | 1 | 0.56 | 0.667 | 0.58 | 0.62 |
| | High | 1 | 1 | 1 | 0.5 | 0.688 | 0.579 | 1 | 0.593 | 0.744 | 0.833 | 0.76 | 0.795 |
| | Micro_average | 0.799 | 0.799 | 0.799 | 0.558 | 0.558 | 0.558 | 0.776 | 0.776 | 0.776 | 0.711 | 0.711 | 0.711 |
| | weighted_average | 0.857 | 0.799 | 0.827 | 0.575 | 0.558 | 0.566 | 0.913 | 0.776 | 0.839 | 0.782 | 0.711 | 0.744 |
| EN_r_a | Low | 0.713 | 1 | 0.832 | 0.576 | 1 | 0.731 | 1 | 0.847 | 0.917 | 0.763 | 0.949 | 0.846 |
| | Medium | 1 | 0.494 | 0.662 | 1 | 0.244 | 0.393 | 0.28 | 1 | 0.438 | 0.76 | 0.58 | 0.658 |
| | High | 1 | 1 | 1 | 0.593 | 1 | 0.744 | 1 | 0.204 | 0.338 | 0.864 | 0.735 | 0.794 |
| | Micro_average | 0.799 | 0.799 | 0.799 | 0.633 | 0.633 | 0.633 | 0.633 | 0.633 | 0.633 | 0.688 | 0.688 | 0.688 |
| | weighted_average | 0.857 | 0.799 | 0.827 | 0.786 | 0.633 | 0.701 | 0.897 | 0.633 | 0.742 | 0.847 | 0.688 | 0.759 |
| EN_m_a | Low | 0.789 | 0.789 | 0.789 | 0.49 | 1 | 0.658 | 1 | 0.847 | 0.917 | 0.76 | 0.879 | 0.815 |
| | Medium | 0.735 | 0.735 | 0.735 | 1 | 0.415 | 0.586 | 0.323 | 1 | 0.488 | 0.686 | 0.717 | 0.701 |
| | High | 1 | 1 | 1 | 1 | 1 | 1 | 1 | 0.389 | 0.56 | 1 | 0.796 | 0.887 |
| | Micro_average | 0.79 | 0.79 | 0.79 | 0.716 | 0.716 | 0.716 | 0.701 | 0.701 | 0.701 | 0.735 | 0.735 | 0.735 |
| | weighted_average | 0.79 | 0.79 | 0.79 | 0.861 | 0.716 | 0.781 | 0.903 | 0.701 | 0.789 | 0.851 | 0.735 | 0.789 |
| V_r | Low | 0.662 | 0.789 | 0.72 | 0.49 | 1 | 0.658 | 1 | 0.847 | 0.917 | 0.717 | 0.879 | 0.79 |
| | Medium | 0.589 | 0.38 | 0.462 | 1 | 0.074 | 0.137 | 0.488 | 1 | 0.656 | 0.692 | 0.484 | 0.57 |
| | High | 0.698 | 1 | 0.822 | 0.593 | 1 | 0.744 | 1 | 0.796 | 0.887 | 0.763 | 0.932 | 0.839 |
| | Micro_average | 0.648 | 0.648 | 0.648 | 0.55 | 0.55 | 0.55 | 0.85 | 0.85 | 0.85 | 0.683 | 0.683 | 0.683 |
| | weighted_average | 0.637 | 0.648 | 0.642 | 0.763 | 0.55 | 0.639 | 0.927 | 0.85 | 0.887 | 0.775 | 0.683 | 0.726 |
| V_b | Low | 0.662 | 0.789 | 0.72 | 0.697 | 1 | 0.822 | 1 | 0.695 | 0.82 | 0.786 | 0.828 | 0.806 |
| | Medium | 0.589 | 0.38 | 0.462 | 1 | 0.415 | 0.586 | 0.244 | 1 | 0.393 | 0.611 | 0.598 | 0.605 |
| | High | 0.698 | 1 | 0.822 | 0.593 | 1 | 0.744 | 1 | 0.204 | 0.338 | 0.763 | 0.735 | 0.749 |
| | Micro_average | 0.648 | 0.648 | 0.648 | 0.716 | 0.716 | 0.716 | 0.558 | 0.558 | 0.558 | 0.641 | 0.641 | 0.641 |
| | weighted_average | 0.637 | 0.648 | 0.642 | 0.819 | 0.716 | 0.764 | 0.892 | 0.558 | 0.687 | 0.783 | 0.641 | 0.705 |
| V_r_a | Low | 0.826 | 1 | 0.904 | 0.616 | 0.697 | 0.654 | 1 | 0.57 | 0.726 | 0.814 | 0.756 | 0.784 |
| | Medium | 1 | 0.62 | 0.766 | 0.816 | 0.756 | 0.785 | 0.221 | 1 | 0.362 | 0.679 | 0.792 | 0.731 |
| | High | 0.698 | 1 | 0.822 | 1 | 1 | 1 | 1 | 0.204 | 0.338 | 0.899 | 0.735 | 0.809 |
| | Micro_average | 0.849 | 0.849 | 0.849 | 0.798 | 0.798 | 0.798 | 0.497 | 0.497 | 0.497 | 0.715 | 0.715 | 0.715 |
| | weighted_average | 0.881 | 0.849 | 0.865 | 0.806 | 0.798 | 0.802 | 0.889 | 0.497 | 0.637 | 0.859 | 0.715 | 0.78 |
| V_m_a | Low | 0.662 | 0.789 | 0.72 | 0.697 | 1 | 0.822 | 1 | 0.264 | 0.418 | 0.786 | 0.684 | 0.732 |
| | Medium | 0.651 | 0.494 | 0.562 | 1 | 0.415 | 0.586 | 0.219 | 1 | 0.359 | 0.623 | 0.636 | 0.63 |
| | High | 1 | 1 | 1 | 0.593 | 1 | 0.744 | 1 | 0.593 | 0.744 | 0.864 | 0.864 | 0.864 |
| | Micro_average | 0.694 | 0.694 | 0.694 | 0.716 | 0.716 | 0.716 | 0.49 | 0.49 | 0.49 | 0.633 | 0.633 | 0.633 |
| | weighted_average | 0.693 | 0.694 | 0.693 | 0.819 | 0.716 | 0.764 | 0.888 | 0.49 | 0.632 | 0.8 | 0.633 | 0.707 |
